# Developmental and housekeeping transcriptional programs display distinct modes of enhancer-enhancer cooperativity in *Drosophila*

**DOI:** 10.1101/2023.10.10.561770

**Authors:** Vincent Loubiere, Bernardo P. de Almeida, Michaela Pagani, Alexander Stark

## Abstract

Genomic enhancers are key transcriptional regulators which, upon the binding of sequence-specific transcription factors, activate their cognate target promoters. Although enhancers have been extensively studied in isolation, a substantial number of genes have more than one simultaneously active enhancer, and it remains unclear how these cooperate to regulate transcription. Using *Drosophila* melanogaster S2 cells as a model, we assay the activities of more than a thousand individual enhancers and a million enhancer pairs towards housekeeping and developmental core promoters with STARR-seq. We report that housekeeping and developmental enhancers show distinct modes of enhancer-enhancer cooperativity: while housekeeping enhancers are additive such that their combined activity mirrors the sum of their individual activities, developmental enhancers are synergistic and follow a multiplicative model of cooperativity. This developmental enhancer synergy is promiscuous and neither depends on the enhancers’ endogenous genomic contexts nor on specific transcription factor motif signatures, but it saturates for the highest levels of enhancer activity. These results have important implications for our understanding of gene-regulation in complex multi-enhancer loci and genomically clustered housekeeping genes, providing a rationale for strong and mild transcriptional effects of mutations within enhancer regions.

## Introduction

A key goal in biology is to understand how gene transcription is regulated, as it represents the first step for a gene to exert its biological function. This task has proven difficult due to the complexity of gene cis-regulatory landscapes, which typically encompass several discrete regulatory elements, termed enhancers, that jointly shape the activity of their target gene’s cognate core promoter^1–3^ (CP) and thus gene transcription. Adding to this complexity, cell-type specific developmental genes and housekeeping genes are regulated *via* two distinct transcriptional programs in *Drosophila* and their transcription relies on different transcription factors^4^ (TFs) and co-factors^5^ (COFs).

In the past years, the question of how several concomitantly active enhancers cooperate to drive transcription received increasing attention^6–8^. In other terms, do different enhancers that are each individually active combine their gene-regulatory functions additively, super-additively (synergistically or multiplicatively) or sub-additively (redundantly) towards their target CP? This question is essential because non-coding mutations affecting synergistic enhancers have an oversized impact on transcription *in situ*, and are therefore more likely to cause downstream functional defects and/or diseases^8^. However, sparse attempts to understand how enhancer-enhancer cooperativity shapes transcription yielded inconsistent outcomes, while multi-enhancer *loci* are common in both flies^1^ and mammals^2^.

Early studies suggested that enhancers are additive^8–12^, meaning that their combined transcriptional outcome mirrors the sum of their individual activities. However, super-additive (or synergistic)^8,10,13,14^ and sub-additive modes^10,15^ have also been reported, whereby combined enhancers are either stronger or weaker than their summed activities. For example, *knirps* enhancers have been shown to exhibit additive or super-additive activities in developing *Drosophila* embryos, while *hunchback* enhancers are sub-additive^10^. In mammals, synergistic enhancers were found to be over-represented at cell-type specific *loci*^7^ and their function was proposed to rely on the formation of co-factor condensates^7^. Nevertheless, these observations were either inferred from a limited number of enhancer combinations or by using correlative strategies that didn’t directly measure the enhancers’ individual and combined activities. As such, the relative proportion of additive *versus* non-additive modes of cooperativity and the gene-regulatory contexts in which they are employed remain unclear, as systematic approaches to quantitatively assess such interactions at high throughput are lacking.

Here, we developed an efficient method to simultaneously assess the activity of many individual enhancers and the corresponding pairwise combinations using a single, internally normalized STARR-Seq assay. Using *Drosophila* S2 cells as a model system, we measured the individual activities of more than a thousand candidate sequences – spanning a wide range of enhancer activities – and more than a million pairwise combinations in a tightly controlled, tractable environment. Our results indicate that developmental enhancers that activate tissue-specific genes are synergistic: the activity of enhancer pairs can be accurately predicted using a simple multiplicative model. Consistently, no specific DNA motif signature was associated with enhancers or enhancer pairs displaying stronger or weaker synergy, arguing against the existence of dedicated synergistic enhancers or enhancer pairs and suggesting a flexible DNA motif syntax supporting promiscuous synergistic interactions. In stark contrast, enhancers that activate housekeeping genes behave additively, i.e. their combined activity corresponds to the sum of the individual activities. This functional difference is associated with higher fraction of Intrinsically Disordered Regions (IDRs) within developmental TFs, which might support downstream synergistic interactions^8,16^, while housekeeping regulation might build on the know propensity of housekeeping genes to cluster along the *Drosophila* genome^17^.

## Results

### High-throughput quantitative assessment of enhancer-enhancer pairs

To tackle the modes of enhancer cooperativity at high-throughput, we developed a new approach to simultaneously measure the activity of many individual enhancers as well as all pairwise combinations in a single, tightly controlled STARR-Seq^1^ assay (Fig. 1a). To achieve this, we designed a pool of 249bp oligos containing 850 enhancers covering a wide range of activities in *Drosophila* S2 cells, together with 150 randomly selected control sequences (see Methods, Supplementary Tables 1-2). We then developed an efficient, fusion PCR-based strategy to systematically fuse these 1,000 sequences to the 5’ and 3’ ends of a transcriptionally inert 300bp spacer sequence, resulting in 1 million combinations including enhancer-enhancer, enhancer-control, control-enhancer and control-control pairs (see Methods, Supplementary Fig. 1a and Supplementary Table 3). Then, resulting constructs were cloned downstream of a developmental CP (dCP), so that the activity of each pair will be reflected by its self-transcription (Fig. 1a). We then followed the UMI-STARR-seq protocol^18^ to perform the functional screens in *Drosophila* S2 cells.

**Figure 1:**
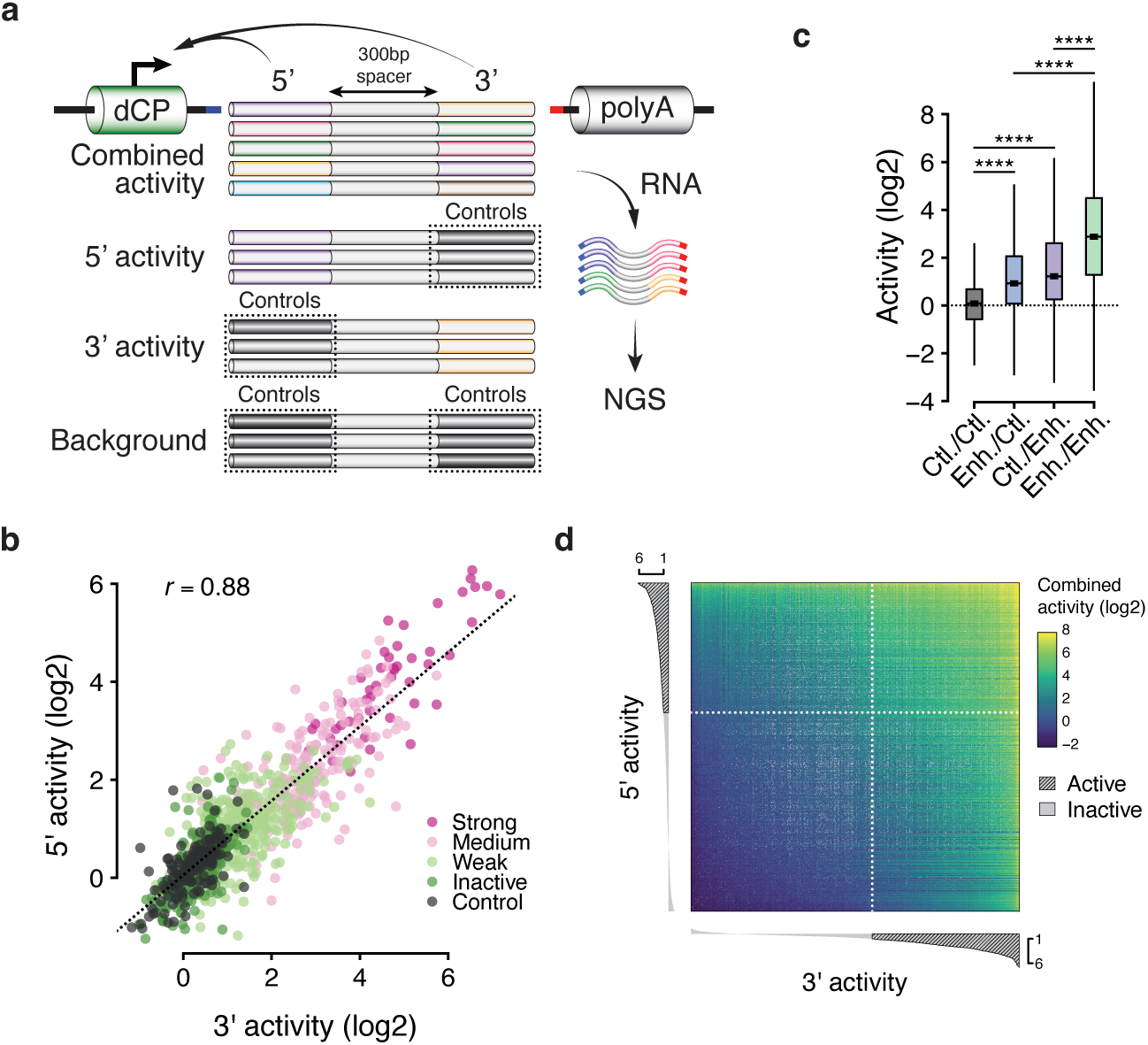
High-throughput assessment of the individual and combined activities of many enhancers. a- Overview of the STARR-Seq reporter assay used to simultaneously measure the individual and combined activities of many enhancers. Random control (in grey) and candidate sequences (colors) are fused to the 5’ and the 3’ ends of a transcriptionally inert spacer and cloned down- stream of a core-promoter, whose transcription mirrors enhancers’ individual and combined activities. b- Correlation between 3’ (x axis) and 5’ (y axis) individual activities of 953 candidate sequences. The dotted line represents the fitted linear model, and color code displays publicly available activity data measured by regular STARR-Seq. Pearson’s correlation coefficient (r) is shown on top left. c- Quantification of the activity of pairs consisting of two random control sequences (Ctl./Ctl., in grey), one control sequence paired with a candidate sequence either in the 5’ (Enh./Ctl., in blue) or the 3’ (Ctl./Enh., in purple) location, or two candidate sequences (Enh./Enh., in green). Wilcoxon test, ****pval<1e-5). d- Heatmap of paired activities (see color legend) ranked by individual activities of the 3’ (x axis) and 5’ (y axis) candidate sequences. 3’ and 5’ activities are depicted as bar charts on x an y axes, respectively, with active sequences being highlight with dashed lines (log2 individual activity >1).

Using this approach, we were able to measure the individual activity of 970 and 961 candidate sequences either in the 5’ and 3’ locations, respectively, and the combined activity of 715,479 pairs (see Methods and Supplementary Table 4). STARR-Seq biological replicates showed a Pearson’s correlation coefficient (*r*) of 0.95 and the inferred activities could be validated quantitatively using luciferase assays (*r*= 0.81), indicating that the method is highly reproducible and robust (Supplementary Fig. 1b-c). Moreover, the individual enhancers’ activities in the 5’ and 3’ locations (inferred using enhancer-control and control-enhancer pairs, respectively, see Methods) were highly correlated (*r*= 0.88) and agreed well with previously published activities STARR-Seq data^19^ (Fig. 1b), indicating that the increased reporter-transcript length and the spacer sequence do not interfere with enhancer function and STARR-seq processing. Furthermore, enhancer pairs were globally stronger than enhancer-control or control-enhancer pairs (Fig. 1c), indicating that two enhancer sequences typically contribute concurrently to transcriptional activation. Accordingly, the combined enhancer activities scaled with the individual enhancer activities, whereby the presence of a single enhancer was sufficient to drive transcription and maximum activities were achieved by combining two strong enhancers (Fig. 1d). Together, these data show the robustness of our method and its unique potential to directly measure the activity of many individual enhancers and enhancer-enhancer pairs at an unprecedented scale.

### Developmental enhancers synergize multiplicatively

To tackle how enhancers cooperate in pairs, we aimed at predicting combined activities using either an additive model – which posits that both candidates work independently from one another – or a multiplicative model, that assumes synergistic interactions between candidates. Importantly, the additive model tended to under-predict the observed activities of enhancer pairs and yielded a rather modest adjusted R-Squared (R^2^) of 0.72 that was substantially outperformed by the multiplicative model (R^2^= 0.83, Fig. 2a). The predictions of the multiplicative model were more accurate for 90% of enhancer-enhancer pairs (in which both candidate sequences are active, Fig. 2b), indicating that developmental enhancers generally synergize.

**Figure 2:**
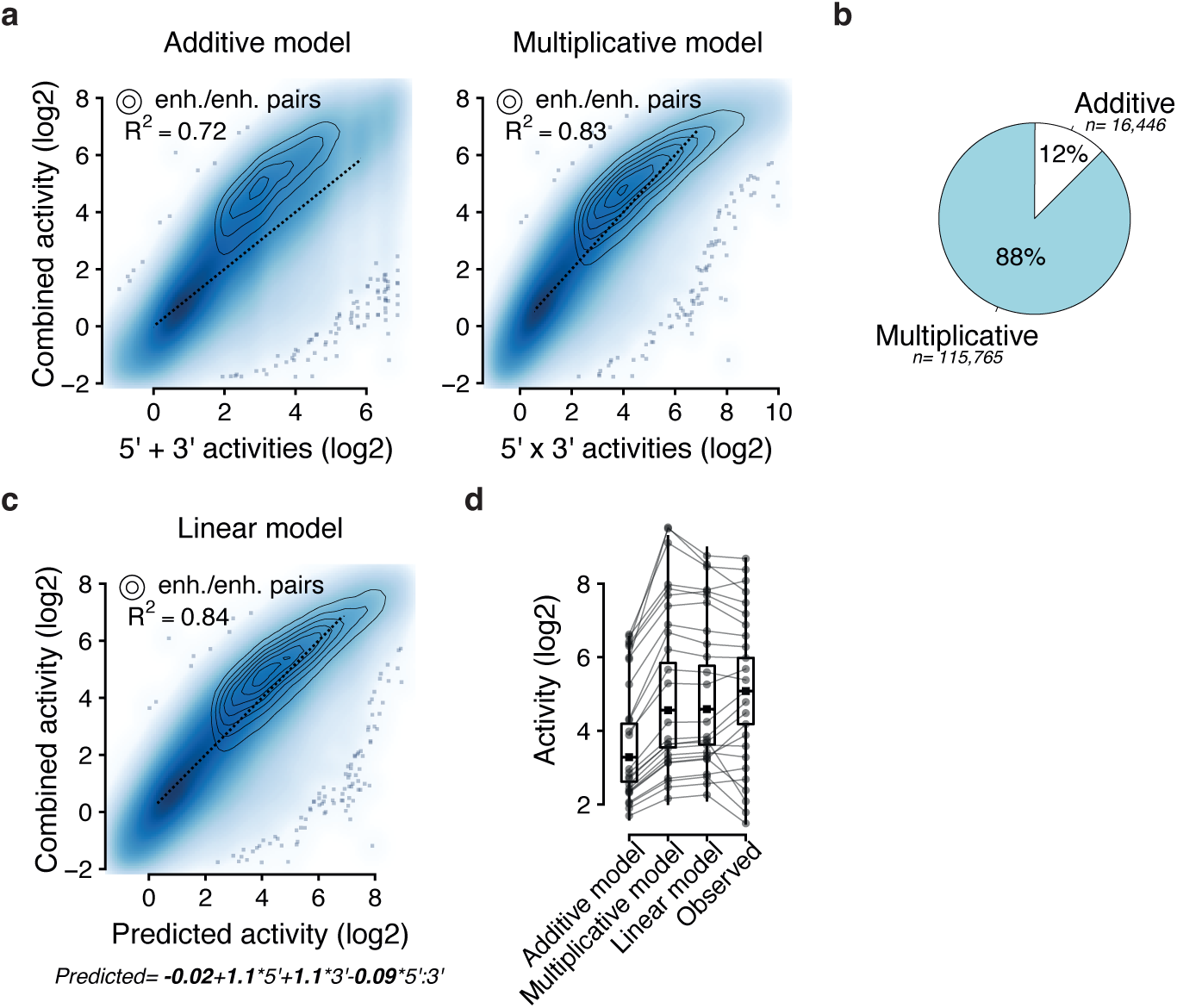
Developmental enhancers synergize multiplicatively. a- Scatterplots showing expected activities (x axis) based on an additive (left) or a multiplicative model (right) versus observed activities (y axis), with corresponding R-squared (R2) values. Enhancer-enhancer pairs in which both candidate sequences are active (see Methods) are highlighted using density lines (see legend) and dotted lines correspond to identity lines (y=x). b- Fraction of enhancer-enhancer pairs (in which both candidate sequences are active) for which the additive (in white) or the multiplicative (in blue) predicted values were the most accurate. Total number of pairs is shown for each category. c- Linear model with interaction using 5’ and 3’ individual activities to predict combined activities, with the corresponding R2 value (top left). Enhanc- er-enhancer pairs in which both candidate sequences are active are highlighted using density lines (see legend). Dotted lines correspond to the identity line (y=x) where expected and observed values are equal. Fitted coefficients and resulting equation are shown on the bottom. d- Boxplot showing, for all enhancer-enhancer pairs, expected (right) and observed activity values (left) using the three different models. Lines connect the values for a representative set of pairs, spanning the whole dynamic range of observed activities.

To further dissect the relationship between individual and combined activities, we fitted a linear regression aiming to predict the activity of enhancer-enhancer pairs by combining the 5’ and 3’ individual activities in a weighted fashion. Of note, the linear regression was fitted using log2 activity values, meaning that fitted coefficients should be interpreted in the context of a multiplicative model (see Methods). The linear model slightly improved the prediction accuracy further (R^2^= 0.84, Fig. 2c-d and Supplementary Fig. 2a-b). One key asset of linear models is to deliver a set of interpretable and informative coefficients. Here, the intercept of −0.02 indicates that pairs consisting of two inactive sequences generally remain inactive, as expected. On the other hand, 5’ and 3’ coefficients (of 1.13 and 1.08, respectively) were both similar and close to 1.0, indicating that optimal predictions were achieved by assuming that both enhancers similarly contribute to the activity of the pair, each to an extent that mirrors its individual activity. Hence, these coefficients substantially agree with a simple multiplicative model.

Interestingly, the linear model revealed a slight yet significant negative interaction between the two enhancers’ activities (coefficient= −0.09), suggesting that the presence of strong enhancers is associated with reduced synergy. This interaction lowers predictions in the highest activity range, for which the multiplicative model tends to over-predict (Supplementary Fig. 2c). This presumably indicates that the CP saturates in the presence of very strong enhancers, a phenomenon which has already been shown to constrain enhancer-promoter function^20^. Consistently, the measured activity of the strongest 5’ or 3’ enhancers can hardly be increased by adding another enhancer within the pair (Supplementary Fig. 2d-e).

Altogether, these data suggest a rather simple view, whereby developmental enhancers would cooperate multiplicatively until saturating their cognate CP.

### Developmental enhancer synergy is promiscuous

The multiplicative linear model can accurately predict the activity of enhancer pairs, using only the activity of individual enhancers and no additional information (regarding for example the enhancers’ sequences or native genomic contexts). Thus, it precludes the existence of large proportions of non-synergistic interactions (e.g., additive or sub-additive) and rather suggests that developmental enhancers are generally synergistic, challenging the existence of complex rules for how they interact with each other. Nevertheless, some pairs remain stronger or weaker than predicted (Fig. 2c, 3a), which could reflect additional rules not considered by the multiplicative linear model or the natural spread of the distribution of synergies. We thus sought to investigate whether these differences – referred to as “residuals” – could be associated with specific DNA motif signatures since, in the STARR-Seq setup, enhancer pairs only differ by their DNA sequences (while everything else is kept constant).

**Figure 3:**
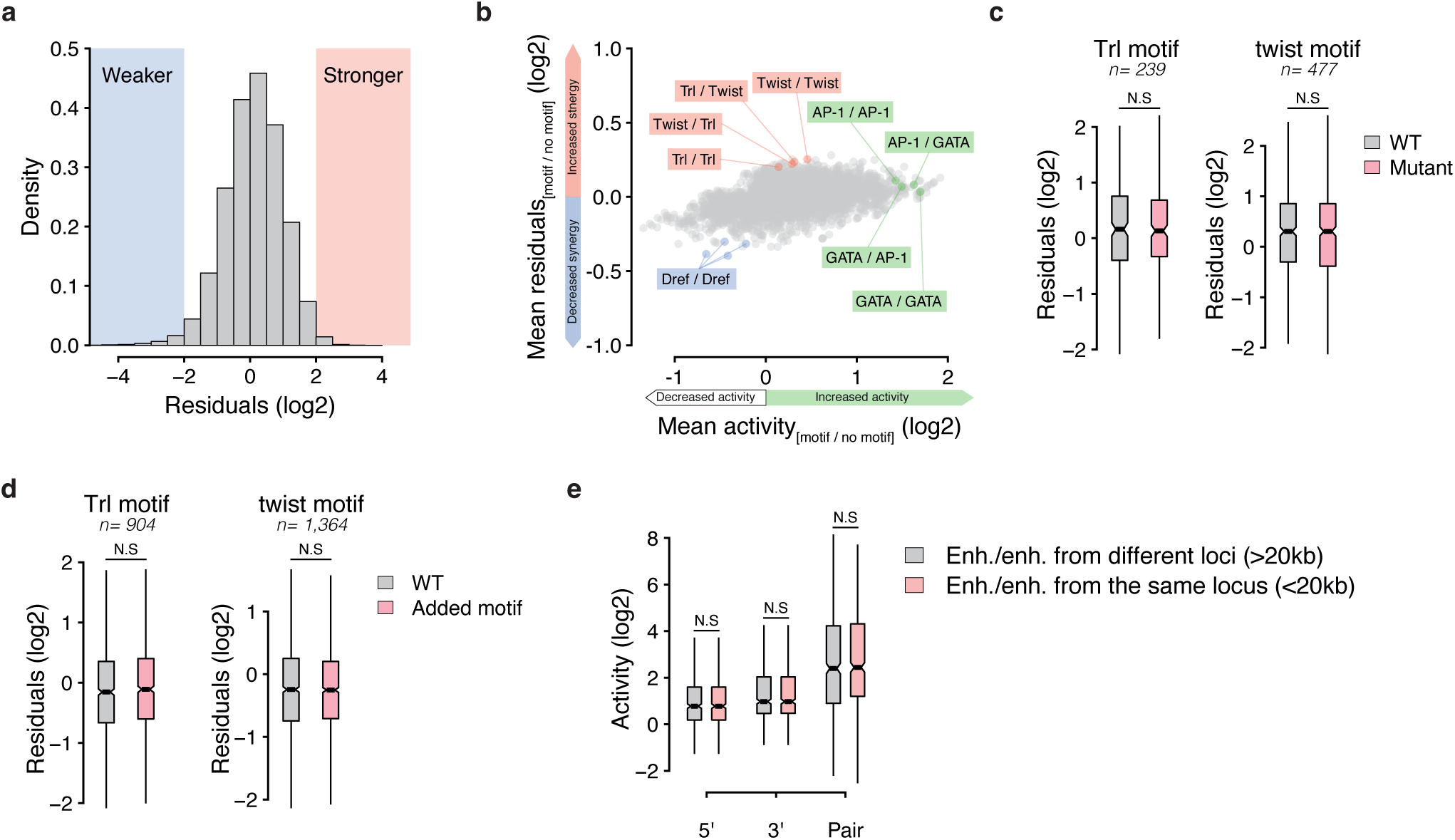
Developmental enhancer synergy is promiscuous. a- Distribution of the residuals of the linear model, ranging from negative (weaker synergy, in blue) to positive (stronger synergy, in orange). b- For each pairwise combination of TF motifs (57*57= 3,249), the overall impact on the activity (x axis) and synergy (y axis) of corresponding enhancer-enhancer pairs are shown (see Methods). For example, Trl/Twist pairs (in which the 5’ and 3’ enhancers contain at least one instance of the Trl and Twist motifs, respectively) show globally increased residuals (in orange). AP-1 and GATA motifs strongly affected enhancer activity while Dref motifs negatively impacted synergy. c-d- Impact of mutating (c, in pink) or adding (d, in pink) Trl (left) or Twist motifs (right) on the synergy of corresponding wild-type (WT) developmental enhancer pairs (in grey), measured as the residuals of the linear multiplicative model (see Methods). e- Quantitative comparison of the activity of enhancer-enhancer pairs from the same locus (<20kb distance in situ, in pink) versus an activity-matched set of distant enhancer-enhancer pairs (in grey, see Methods). Wilcoxon test, N.S= Not Significant, ***=pval<1e-3.

We therefore systematically measured the association of a large collection of TF motifs with the enhancer activities and with the residuals, the latter ones reflecting the difference between expected synergistic and observed activities (see Methods). In line with their prominent role in driving enhancer activity in S2 cells^19,21^, homotypic and heterotypic combinations of the AP-1 and GATA motifs (either in the 5’ or 3’ enhancers) were associated with stronger overall enhancer activity, but not with increased or decreased synergy (Fig. 3b). Compared to the range of activities (from −1 to +2 on a log2 scale), the range of residuals is substantially smaller (comprised between −0.5 and +0.5, approximately, Fig. 3b). This is consistent with the accuracy of the activity-based multiplicative model and argues that developmental enhancer synergy is promiscuous in S2 cells, with additional rules having only minor influences.

To nevertheless test whether enhancer synergy might be influenced by specific TF motifs, we decided to focus on the Trl motif (GAGA) and a variant of the Twist motif (CATATG), which were associated with high residuals but not with enhancer activity overall (Fig. 3b). We selected 50 enhancers containing each of these motifs (see PWMs in Supplementary Table 5), mutated them and measured the activity of the resulting pairs using STARR-Seq (Supplementary Tables 6-7). Based on the fitted multiplicative model, wild-type (wt) and mutated pairs showed similar residuals (Fig. 3c). Similarly, pasting these motifs into inactive control sequences or into enhancers that did not contain them also had no substantial impact on synergy (Fig. 3d), suggesting that developmental enhancer synergy does not strongly rely on these two motifs. Consistently, a LASSO regression using motif counts as input was able to predict the overall activity of enhancer pairs (R^2^= 0.37) but not the residuals (R^2^ = 0.08, Supplementary Fig. 3a-b), confirming the association between TF motifs and enhancer activities and suggesting that residuals or synergies do not rely on specific DNA binding motifs. Overall, our results indicate that synergistic interactions between developmental enhancers do not rely on rigid motif syntax rules nor specific combinations of TFs, and might rather be promiscuous in our system. In line with this idea, pairs consisting of two enhancers closely spaced in their endogenous loci (natively less than 20kb apart), which might have evolved to cooperate more efficiently, are not notably stronger than control pairs of distant enhancers (natively more than 20kb, see Fig. 3e and Methods).

### Housekeeping enhancers are additive

In *Drosophila*, tissue-specific developmental genes and housekeeping genes form two different transcriptional programs which rely on distinct set of enhancers, CPs and TFs^4,5^. Dref is a key regulator of housekeeping genes in *Drosophila*^5,22^ and, interestingly, its DNA binding motif was associated with substantially lower residuals (Fig. 3b), suggesting that it might impair synergistic interactions between developmental enhancers. Consistently, pasting Dref motifs within developmental enhancers significantly reduced their residuals but also their individual activities (Supplementary Fig, 3b-c). Hence, it is unclear whether this reduced synergy is related to the developmental CP, which cannot be efficiently activated by housekeeping-type enhancers and TFs.

To address this question, we decided to assess the activities of enhancers and enhancer pairs towards the RpS12 housekeeping CP. We designed a smaller synthetic DNA library containing 62 housekeeping enhancers, 53 developmental enhancers and 50 control sequences (Supplementary Table 8), paired all sequences systematically as above, and cloned all pairs downstream of the RpS12 housekeeping CP. We then performed STARR-Seq to measure the activities of the individual enhancers and enhancer pairs as described and modeled the data with the additive and multiplicative models (see Supplementary Table 9). Contrasting with the previous developmental setup, the additive model could accurately predict the activity of housekeeping enhancer-enhancer pairs (R^2^= 0.78) and significantly outperformed the multiplicative model (R^2^= 0.19, Fig. 4a-b). Hence, housekeeping enhancers interact additively, even in the presence of a compatible CP.

**Figure 4:**
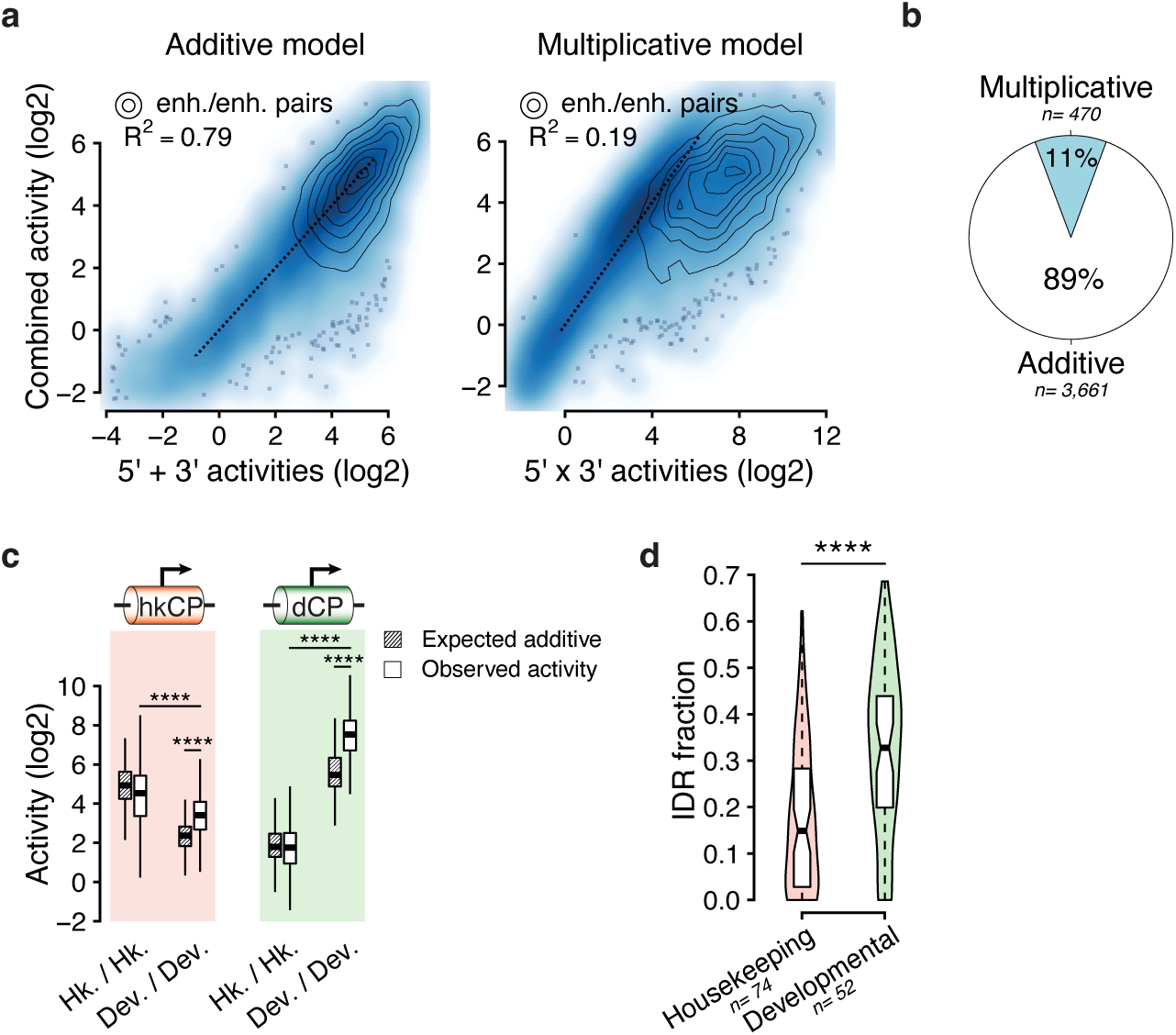
Housekeeping enhancers are additive and synergistic versus additive modes of cooperativity do not depend on the CP type. a- Scatterplots showing expected activities (x axis) based on an additive (left) or a multiplicative model (right) versus observed activities (y axis) using the RpS12 housekeeping CP (on top). Corresponding R-squared (R2) values are shown (top left) and enhancer-enhancer pairs in which both candidate sequences are active are highlighted using density lines (see legend). Identity lines (where expected and observed values are equal) are shown using dotted lines. b- Fraction of enhancer-enhancer pairs (in which both candidate sequences are active) for which the additive (in white) or the multiplicative (in blue) predicted values were the most accurate. c- Expected additive and observed activities of housekeeping versus developmental enhancer-enhancer pairs (x axis) using either and housekeeping (hkCP, in red) or a developmental (dCP, in green) core-promoter (on top). Wilcoxon test, ****=pval<1e5. d- Fraction of Intrinsically Disordered Regions (IDRs) within TF and COFs proteins showing a preference in activating Housekeeping (Hk., in orange) versus Developmental (Dev., in green) CPs (see Methods). Wilcoxon test, ****=pval<1e-5.

To further compare developmental and housekeeping contexts side-by-side, we also measured the activity of this smaller library using the DSCP developmental core-promoter (see Supplementary Table 10). First, we note that the known specificity between developmental and housekeeping enhancer-CP^4^ holds true: with the housekeeping CP, maximum activities are achieved by combining two housekeeping enhancers, while the developmental CP reaches its maximum levels with developmental enhancer pairs (Fig. 4c). Importantly however, the developmental enhancers are synergistic and the housekeeping enhancers additive irrespective of CP type, indicating that synergistic versus additive modes of cooperativity are enhancer-intrinsic properties independent of the CP.

Together, our results indicate that housekeeping and developmental enhancers are intrinsically different in their modes of cooperativity, namely additive versus synergistic, respectively, and that this inherent distinction is independent of their interaction with the CP.

## Discussion

Here, we developed a new approach to study enhancer-enhancer cooperativity at an unprecedented scale, uncovering an unexpected discrepancy between developmental and housekeeping transcriptional programs in *Drosophila*: while developmental enhancers activating tissue-specific genes interact synergistically, housekeeping enhancers behave additively.

Further dissection of the developmental dataset suggests a rather simple model, where the activity of the two enhancers combine multiplicatively, until they eventually saturate the CP. Thus, synergistic interactions between regulatory elements might be more widespread than previously thought, with a recent study showing that enhancer and promoter activities multiplicatively combine to determine RNA output in mammals^20^. Widespread synergistic cooperativity has important implications, since a single mutation affecting a synergistic enhancer might have a drastic effect on transcription^10^ and potentially influence disease risk ^8^. On the other hand, enhancer synergy plus promoter saturation might also foster the known robustness of developmental *loci* containing many enhancers^2^: given our finding that CP saturation constrains the transcriptional outcome of pairs of very strong enhancers, even the full mutational disruption of one enhancer would have no impact as the remaining enhancers would be sufficient to drive maximal transcriptional outcome. At such complex *loci*, removing one or several enhancers has indeed no impact on transcription^2^ (i.e. enhancers act redundantly), implying that the combined activity of enhancers initially exceeds the capacity of the promoter.

Interestingly, we did not find specific DNA motif signatures that would support selectively enhanced synergy, arguing against the existence of strong specificities between subsets of developmental enhancers. Thus, the way TFs and COFs interact to dictate transcription might slightly differ when considering enhancer pairs *versus* single enhancers. In a single developmental enhancer, distinct combinations of TFs were shown to exhibit distinct types of additive and cooperative behaviors^19,23,24^. In contrast, our results suggest that higher order interactions between the ensemble of TFs and COFs that each developmental enhancer can recruit are less specific, since they globally lead to synergistic interactions. However, the *Drosophila* genomic enhancer sequences we have been using typically already contain homotypic and heterotypic combinations of motifs, and future studies could use synthetic sequences to more specifically assess the impact of each motif.

Contrasting with developmental enhancers, we found housekeeping enhancers to behave additively, suggesting that the molecular mechanisms governing the transcription of *Drosophila* developmental and housekeeping genes are fundamentally different. Notably, a recent study suggested that synergistic interactions might indeed be characteristic of developmental *loci* in mouse, while other enhancers would be additive^7^. Altogether, these findings lead us to postulate that developmental TFs might have specifically evolved the ability to support synergistic interactions in order to support sharp transcriptional changes in response to developmental cues. In recent years, extensive investigations have focused on the function of Intrinsically Disordered Domains (IDRs), whose presence in TFs/COFs has been repeatedly associated with the formation of condensates^16^ and synergistic transcriptional outcomes^8^. By comparing TFs and COFs showing a preference for either developmental or housekeeping enhancers^25^ (see Methods), we interestingly found that developmental TFs/COFs contain a significantly higher fraction of IDRs (32.8% vs. 14.9% on average, Fig. 4d), which might favor synergistic interactions at developmental enhancers in *Drosophila*. However, further studies would be needed to tackle how additive *versus* synergistic behaviors are encoded at the protein level.

Finally, additive interactions seem sufficient to foster steady transcription of housekeeping genes. However, such interactions still imply that housekeeping enhancers might boost each, which could explain why housekeeping genes and enhancers tend to form clusters along the *Drosophila* genome, an arrangement that has previously been shown to be important for their proper transcription^17^.

**Supplementary Figure 1:**
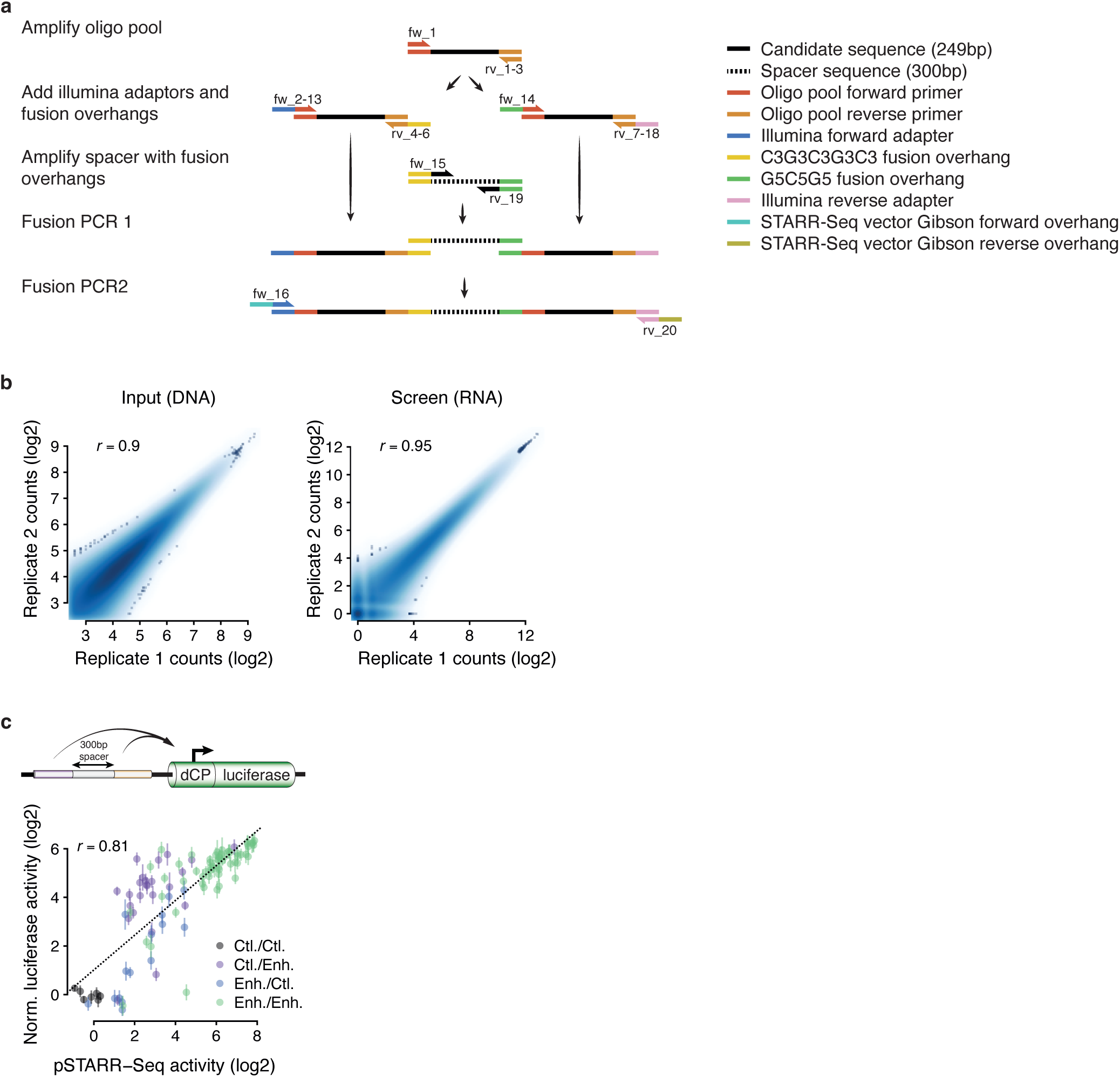
a- Schematic view of the fusion PCR protocol used to generate complex pools of enhancer-enhancer pairs. b- Correlation between RNA (screen) and DNA (input) counts between two STARR-Seq replicates. Pearson’s correlation coefficient (r) is shown (top left). c- Correlation between STARR-Seq (x axis) and luciferase (y axis) measurements for a set of control-control random sequences (Ctl./Ctl., in grey), one control sequence paired with a candidate sequence either in the 5’ (Enh./Ctl., in blue) or the 3’ (Ctl./Enh., in purple) location, or two candidate sequences (Enh./Enh., in green). A schematic view of the reporter construct is shown (top), as well as the pearson’s correlation coefficient (r, on the top left of the scatterplot).

**Supplementary Figure 2:**
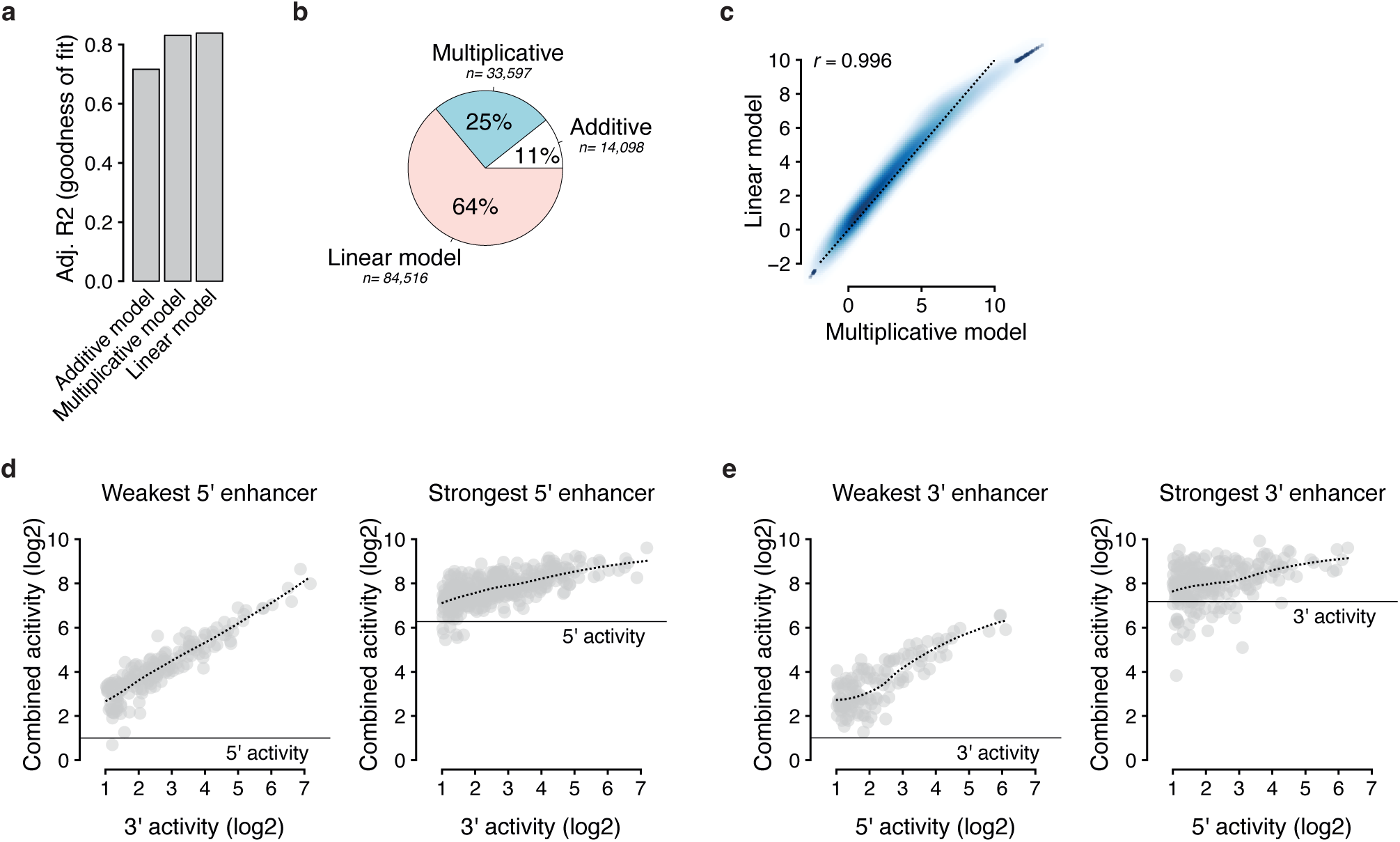
a- Adjusted R-squared (R2) values for the 3 different models (bottom). Higher R2 means better fit. b- Fraction of enhancer-enhancer pairs (in which both candidate sequences are active, see Methods) for which the additive (in white), the multiplicative (in blue) or the multiplicative linear model (in pink) were the most accurate at predicting observed activities. Total number of pairs is shown for each category. c- Scatterplot comparing the predicted values of the multiplicative model (x axis) versus the predicted values of the fitted linear model (y axis). The identify line is shown with a dotted line. d- Scatterplot showing, for all the pairs containing either the weakest (left) or the strongest (right) enhancer in the 5’ location, the relationship between the activity of the other (3’) enhancer and the activity of the pair. While the activity of the weakest 5’ enhancer can be strongly boosted by the other 3’ enhancer, the activity of pairs containing the strongest 5’ enhancer plateau around 8. e- Same as d but considering the weakest (left) and the strongest (right) enhancers being located in the 3’ location.

**Supplementary Figure 3:**
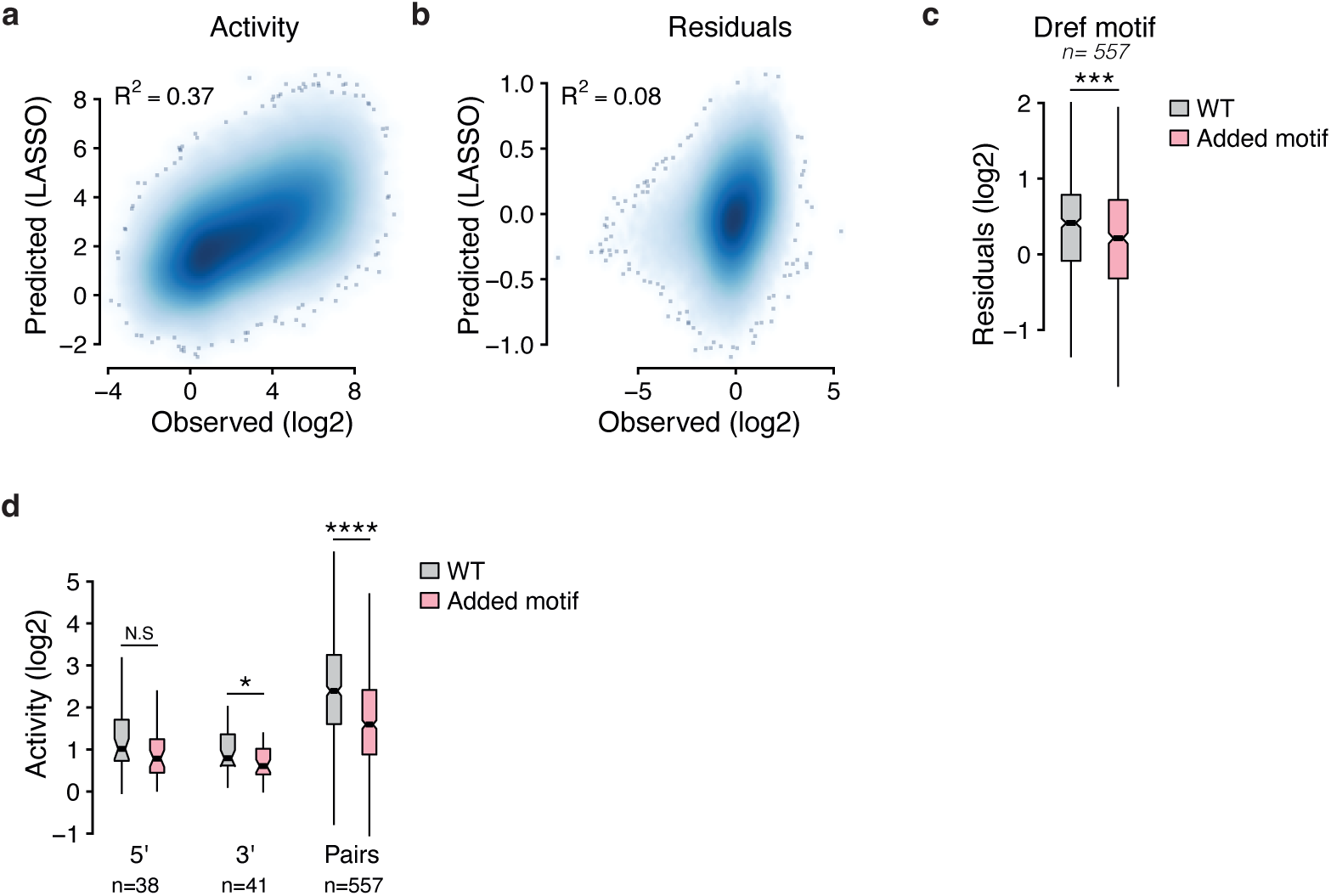
a- Observed enhancer-enhancer pair activities (x axis) versus LASSO predictions (y axis), using a representative set of Drosophila DNA binding motifs. R-squared (R2) value is shown on the top left. b- Observed enhancer-enhancer pair residuals (using the fitted multiplicative model, x axis) versus LASSO predictions (y axis), using a representative set of Drosophila DNA binding motifs. R-squared (R2) value is shown on the top left. c- Impact of adding Dref motifs (in pink) on the synergy of corresponding wild-type (WT) developmental enhancer pairs (in grey), measured as the residuals of the linear multiplicative model (see Methods). c- Impact of adding Dref motifs (in pink) on the individual and combined activities of developmental enhancers (see Methods).

## Supplementary Table legends

**Supplementary Table 1: Oligo pools summary**

For each pool, the number of sequences is reported for each group.

**Supplementary Table 2: WT oligo pool sequences**

IDs and sequences of the 1,000 oligonucleotide sequences of the WT oligo pool. Flanking PCR primers are specified.

**Supplementary Table 3: PCR primers**

PCR primers used to generate enhancer-enhancer pairs.

**Supplementary Table 4: WT oligo pool STARR-Seq activities**

Individual (Ind) and combined STARR-Seq activities (log2FoldChange) of the 5’ (L for “Left”) and 3’ (R for “Right”) measured using the developmental DSCP core promoter and the WT oligo pool.

**Supplementary Table 5: Twist Trl Dref motifs PWMs**

For each pool, the number of sequences is reported for each category.

**Supplementary Table 6: Mutated oligo pool sequences**

IDs and sequences of the 998 oligonucleotide sequences of the Mutated oligo pool. Flanking PCR primers are specified.

**Supplementary Table 7: Mutated oligo pool STARR-Seq activities**

Individual (Ind) and combined STARR-Seq activities (log2FoldChange) of the 5’ (L for “Left”) and 3’ (R for “Right”) measured using the developmental DSCP core promoter and the Mutated oligo pool.

**Supplementary Table 8: Focused oligo pool sequences**

IDs and sequences of the 174 oligonucleotide sequences of the Focused oligo pool. Flanking PCR primers are specified.

**Supplementary Table 9: Focused oligo pool hkCP STARR-Seq activities**

Individual (Ind) and combined STARR-Seq activities (log2FoldChange) of the 5’ (L for “Left”) and 3’ (R for “Right”) measured using the housekeeping RpS12 core promoter (hkCP) and the Focused oligo pool.

**Supplementary Table 10: Focused oligo pool dCP STARR-Seq activities**

Individual (Ind) and combined STARR-Seq activities (log2FoldChange) of the 5’ (L for “Left”) and 3’ (R for “Right”) measured using the developmental DSCP core promoter (dCP) and the Focused oligo pool.

## Material and methods

### Design of oligo pools

For this study, we designed three pools of oligonucleotides, consisting of 249-bp candidate sequences flanked by PCR primers, for a total length of 300nt. For their design, we used publicly available STARR-Seq data^4,19,26^ and DHS data^1^ from *Drosophila* S2 and OSC cells (see the “Data availability” section for corresponding GEO repositories). Unless explicitly mentioned, genomic sequences originated from the dm3 version of the *Drosophila* genome. A summary of the composition of each library is available in Supplementary Table 1. The genomic coordinates and DNA sequences of all oligo pools are available in Supplementary Tables 2,6,8.

A first pool of 1,000 oligos was designed to comprehensively assess how enhancer-enhancer pairs function downstream of a developmental CP. It contained 600 developmental enhancers, 100 housekeeping enhancers, 150 control sequences and 150 inducible/OSC-specific enhancers^4,26^(see “WT oligo pool”, Supplementary Tables 1 and 2). Developmental and housekeeping enhancers were selected to cover a wide range of activities in *Drosophila* S2 cells. For control sequences, we randomly sampled 100 genomic sequences showing no STARR-Seq signal in S2 cells, 25 exon sequences and 25 sequences from 20080805 version of the *E. coli* genome.

To test the impact of Trl, Twist and Dref motifs on cooperativity, we designed a pool of 465 WT sequences and 533 mutated variants (998 in total). WT sequences contained 93 randomly sampled inactive genomic sequences showing no STARR-Seq signal in S2 cells, 106 DHS sites showing no STARR-Seq signal in S2 cells, 131 developmental enhancers containing Twist/Trl motifs, 50 developmental enhancers containing at least two Twist motifs, 50 developmental enhancers containing at least three Trl motifs and 35 enhancers showing both developmental and housekeeping activities (termed “shared” enhancers) containing at least two Dref motifs (see the “Mutated oligo pool” in Supplementary Table 1). The position weight matrices used to identify instances of the Twist/Trl/Dref motifs are available in Supplementary table 5. To assess the relevance of Twist/Trl/Dref motifs, we pasted two or three instances of these motifs into inactive control sequences, DHS sites with no STARR-Seq activity or active developmental enhancers, all of which did not contain endogenous Twist/Trl motifs (398 variants in total). Conversely, we mutated Twist, Trl and Dref motifs by replacing them with random stretches of nucleotides within a set of active enhancers that contained them (135 variants in total). To minimize the impact of pasting/mutating motifs on the individual activity of candidate sequences, we started from a larger pool of WT sequences and generated 1,000 possible variants for each condition, changing the position of pasted motifs and/or the random stretch of nucleotide being used to replace the endogenous motifs. Then, we predicted the activity of all variants using DeepSTARR^19^ and only retained the ones with the smallest impact on predicted individual activities (see Supplementary Table 6).

For the side-by-side comparison of housekeeping and developmental enhancer-enhancer pairs with different CPs, we designed a pool containing 165 candidate sequences: 50 randomly samples inactive sequences showing no STARR-Seq signal, 62 housekeeping and 53 developmental enhancers and 9 shared enhancers, which can activate both developmental and housekeeping CPs (see “Focused oligo pool”, Supplementary Tables 1 and 8).

### Synthesis of enhancer-enhancer pairs

300-mer oligo pools were synthesized by Twist Bioscience, amplified (for PCR primers, see Supplementary Table 3) and split into two batches that will be processed in parallel to generate 5’ and 3’ candidate sequences (see Supplementary Figure 1a for a schematic view of the method). Using overhang PCR, Illumina adapters and CG-rich overhangs were added to the ends of 5’ and 3’ candidates, while the transcriptionally inert spacer was amplified from *Drosophila* genomic DNA (dm3 genomic coordinates: chr2L:509283-509549) and flanked with complementary CG-rich overhangs (primers available in Supplementary Table 3).

This way, the three DNA fragments (5’ candidate, spacer, 3’ candidate) contain overlapping sequences at their extremities, corresponding to CG-rich overhangs that were optimized to allow efficient, orientation-specific fusion PCR reactions^27^ (see Supplementary Figure 1a). Following 20 cycles of linear amplification, fused fragments were amplified for 20 PCR cycles and flanked with overhang sequences compatible with Gibson Assembly® (see primers in Supplementary Table 3). Finally, fused enhancer pairs were size selected using gel purification (~1kb).

### STARR-Seq library cloning and sequencing

STARR-Seq libraries were generated by cloning enhancer-enhancer pairs into the *Drosophila* STARR-Seq vectors, containing either the developmental *Drosophila* synthetic core-promoter^28^ (DSCP) or the housekeeping RpS12 core-promoter^4^. We followed the previously established UMI-STARR-Seq library cloning protocol^18^, except that the In-Fusion HD reaction was replaced by Gibson assembly® (New England BioLabs, #E2611S). Briefly, two Gibson Assembly® reactions were used for each library (500ng of digested plasmid, three molar excess of enhancer-enhancer pair constructs and 40μL 2X Gibson Master Mix, for a total volume of 80μL per reaction) following manufacturer’s instructions. Assembled libraries were electroporated into competent *E. coli* (Invitrogen, #C640003) and grown O/N in 4L LB-Amp (Luria-Bertani medium plus ampicillin, 100 µg/ml) and purified using Plasmid Plus Giga Kit (Qiagen, #12991). Finally, libraries were UMI-tagged and amplified as previously described^18^, and sequenced at the VBCF NGS facility using Next-generation Illumina sequencing, following the manufacturer’s protocol with standard Illumina i5 indexes and UMIs at the i7 index (paired-end 36nt or longer).

### STARR-Seq screens

*Drosophila* S2 cells were cultured at 27°C in Schneider’s *Drosophila* Medium (SM, Gibco, #21720-024) supplemented with 10% heat inactivated FBS (Sigma, #F7524) and passaged every 2-3 days. For each biological replicate, 10^8 cells were resuspended in 400µL of a 1:1 dilution of HyClone MaxCyte electroporation buffer and serum-free SM, and electroporated with 20µg of the input libraries (see previous section) using the MaxCyte-STX system (‘Optimization 1’ protocol). Then, we adapted the established UMI-STARR-Seq protocol^18^ to make it compatible with longer inserts. Sequencing-ready libraries were size selected on a 1% agarose gel (~1kb fragments) and sequenced at the VBCF NGS facility using Next-generation Illumina sequencing, following the manufacturer’s protocol with standard Illumina i5 indexes and UMIs at the i7 index (paired-end 36nt or longer).

### Next generation sequencing data processing

For each oligo pool, a custom indexe containing the corresponding sequences was generated using the buildindex function from the Rsubread R package^29^ (version 2.12.2). STARR-Seq paired-end reads were then aligned using the align function from the same package, with the following parameters: type= “dna”, maxMismatches= 3, unique= TRUE. Only the pairs for which both reads could be aligned with expected orientations and positions were considered. Then, UMI sequences were retrieved, reads were collapsed as previously described^19^, and only the pairs with at least 5 reads in each of the two input replicates were considered for downstream analyses.

### Computation of individual and combined activities

To calculate the activity of all pairs, we added one pseudocount read to UMI-collapsed counts and computed the log2 fold-change over input using DESeq2^30^ (with at least two replicates). To normalize the samples between them and facilitate comparisons between different screens, we used the negative control sequence pair counts as scaling factors, so that the activities of negative control pairs are centered on zero.

Before computing the individual activity of each candidate sequence, we first aimed at removing potential outlier negative control sequence, that might eventually show some activity in our screens. To do so, we assessed the activity of each negative control sequence by averaging its activity across all its observed combinations with other control sequences. Resulting activities were scale in R and only the 5’ and 3’ control sequences with a z-score value located between −1 and 1 were considered as valid, robust control sequences. Finally, these robust control sequences were used to compute the individual activity of each individual candidate sequence (which we refer to as “enhancers” for simplicity), by averaging their activities across all its observed combinations with at least 10 robust control sequences (otherwise, the individual candidate sequence was discarded). To classify an enhancer as active, we compared the activities of all its observed combinations with robust control sequences to the activities of control/control pairs using one-tailed Fisher’s exact tests (alternative= “greater”) followed by false discovery rate (FDR) multiple testing correction. Only the enhancer sequences with a log2 activity bigger than 1 and an FDR<0.05 were considered as active. For each STARR-Seq screen, individual and combined activities are reports in Supplementary Tables 4,7,9,10.

### Modelling of combined activities using individual activities

The individual activity of the 5’ and 3’ enhancers were scaled using negative control sequence pairs. Therefore, they correspond to fold-changes normalized to the basal activity of the core promoter expressed in log2. Hence, for a given pair *P*, log2 additive and multiplicative predicted values were computed using the following formulas:

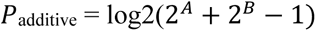

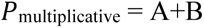

Where *A* and *B* correspond to the log2 individual activities of the 5’ and 3’ enhancers, respectively. To optimize the performance of the multiplicative model, we fitted the following linear model using log2 activity values and the *lm* function in R:

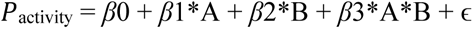

Where *β*0 is the intercept, *β*1 and *β*2 represent the contribution of each enhancer’s activity, *β*3 captures the interaction between the two enhancers and ɛ is the error term. The performance of each model was assessed using adjusted R-squared (R^2^) coefficients.

### DNA binding motifs analyses

To assess the impact of DNA binding motifs on the activity of developmental enhancers and/or synergistic interaction between them, we started from a publicly available collection of 13,899 annotated position weight matrices (PWMs) classified in 901 curated, non-redundant clusters^19,31^. Using publicly available RNA-Seq data^1^, we selected the clusters for which at least one of related PWMs are associated to a TF that is expressed in *Drosophila* S2 cells, resulting in 3,083 motifs from 204 clusters. For each PWM, we counted the number of motifs within all the oligonucleotides of the WT library pool (see previous sections, Supplementary Table 2) and a control set of 1,000 249-bp control sequences (randomly sampled from the dm3 version of the *Drosophila* genome), using the motif_counts function from the motifmatchr R package^32^ (version 1.18.0) with the following parameters: genome= “dm3”, bg = “genome”, p.cutoff = 5e-04. Using two-tailed fisher tests followed by FDR multiple testing correction, we selected the motifs that were significantly over-represented in any of the groups of candidate sequences from the WT oligo pool (see the “group” column for the WT oligo pool in Supplementary Table 1) compared to the set of control random sequences (log2OR>0, FDR < 1e-5) and had at least 10 counts in total; resulting in a curated, non-redundant set of 57 binding motifs.

To assess the effect of a specific combination of motifs on the activity of developmental enhancer-enhancer pairs, we first identified the pairs in which the 5’ and the 3’ enhancers respectively contained the motifs of interest or not. For each combination of motifs (57*57= 3,249), we then computed the mean activity of the pairs that contained the motifs and normalized it to the activity of pairs that did not contain them. We also used this approach to systematically assess the impact of motifs on developmental enhancer synergy, using the mean residuals of the fitted linear model (see previous section), which reflect the difference between predicted multiplicative versus observed values.

Finally, to unbiasedly assess whether DNA binding motifs might be predictive of the activity of an enhancer pair, we trained a LASSO regression to predict the activity of enhancer-enhancer pairs using only motif counts (and no other information). In parallel, we used a similar model to predict linear model’s residuals (see previous section), to assess whether specific combinations of motifs might boost or dampen synergistic interactions between developmental enhancer pairs. For both approaches, we used the glmnet R package^33,34^ (version 4.1-4). The cv.glmnet (alpha = 1, lambda = 10^seq(2, −3, by = −.1), standardize = TRUE, nfolds = 5) function was used to infer the best lambda (bl) value to be passed to the glmnet (alpha = 1, lambda = bl, standardize = TRUE) to function and fit the model.

### IDR content analysis

We used publicly available data reporting the activity of 812 *Drosophila* factors in 24 different enhancer contexts in S2 cells^25^ to identify TFs and COFs that were sufficient to activate either the DSCP (developmental) or the RpS12 (housekeeping) core promoters, respectively (FoldChange>log2(1.5)). Then, TFs/COFs were classified based on their preference for either the housekeeping (RpS12 FoldChange > DSCP FoldChange) or the developmental (RpS12 FoldChange < DSCP FoldChange) core promoter, that are expressed in S2 cells^1^ (rpkm>0.1). This approached identified 74 and 52 TFs/COFs showing a preference for housekeeping and developmental contexts, respectively. For each of them, the fraction of predicted Intrinsically Disordered Regions (IDRs) were retrieved from the MobiDB database^35^.

### Downstream analyses

All downstream analyses were performed in R^36^ (version 4.2.0) using the data.table package^37^ (version 1.14.6). All custom scripts generated for this study are publicly available on GitHub: https://github.com/vloubiere/git_peSTARRSeq.

### Luciferase assays

The promoter of the pGL3 Luciferase Reporter Vector (Promega) was replaced with the DSCP core promoter^4^, and the resulting construct was used to validate pSTARR-Seq measurements. A set of control-control, enhancer-control, control-enhancer and enhancer-enhancer pairs were amplified and flanked with overhangs compatible with Gibson assembly® (5’ overhang= ATTTCTCTATCGATAGGTAC. 3’ overhang= GTACCGAGCTCTTACGCGTC). Resulting sequences were cloned upstream of the core promoter (using KpnI restriction site) *via* Gibson assembly® (New England BioLabs, #E2611S). Each construct was verified using sanger sequencing and luciferase assays were performed as previously described^1^. For each replicate, 6.10^5 S2 cells were transfected with a mix consisting of 100ng of luciferase plasmid and 5ng of Renilla plasmid per well using FuGENE® HD (Promega, #E2311). At least tree biological l replicates were measured per construct, each consisting of 4 technical replicates.

## Data availability

Raw sequencing data and processed files generated for this study will be made public available on GEO prior to publication. Publicly available STARR-Seq data from *Drosophila* S2 and OSC cells were obtained from GSE183939^19^, GSE47691^26^, GSE57876^4^. Publicly available DHS and RNA-Seq data from *Drosophila* S2 cells were obtained from GSE40739^1^.

## Code availability

All custom scripts to analyze the data and plot the figures will be made publicly available on GitHub prior to publication.

## Acknowledgements

We thank the IMP/IMBA/GMI and Max Perutz Labs core facilities for support. Next-generation sequencing was done at the Vienna Biocenter Core Facilities GmbH (VBCF) Next-Generation Sequencing Unit. Research in the Stark group is supported by the Austrian Science Fund (FWF, P29613–B28 and P33157–B). Vincent Loubiere was supported by HFSP (LT000926/2020) and EMBO (790-2019) postdoctoral fellowships. Basic research at the IMP is supported by Boehringer Ingelheim GmbH and the Austrian Research Promotion Agency (FFG). For the purpose of Open Access, the authors have applied a CC BY public copyright license to any Author Accepted Manuscript (AAM) version arising from this submission.

## Author contributions

V.L. and A.S designed the experiments. V.L. developed efficient fusion PCR protocols to generate enhancer-enhancer pairs libraries and performed STARR-Seq experiments together with M.P. V.L. performed bioinformatic analyses with the help of B.P.A. V.L. and A.S. wrote the manuscript, which was read and critically assessed by all authors.

